# Endophyte *Chaetomium globosum* D38 and its elicitors promote tanshinones accumulation of *Salvia miltiorrhiza*

**DOI:** 10.1101/167007

**Authors:** Xin Zhai, Dong Luo, Xiuqing Li, Ting Han, Zhouyang Kong, Jiachen Ji, Luping Qin, Chengjian Zheng

## Abstract

Due to the low yield of tanshinones and their analogues in *Salvia miltiorrhiza*, there are all kinds of stimulation strategies having been applied to improve tanshinones output in plant tissue cultures. Endophytic fungi have formed various different relationships with their host plants withstanding host and environmental factors, including symbiotic, mutualistic, commensalistic, and parasitic. Thus we take the assumption that endophytic fungi may be an emerging microbial tool used to promote secondary metabolism, which will promote the production of active compounds through endophyte-based biology method. Our study therefore aimed to examine the effects of live endophytic fungus *Chaetomium globosum* D38 and its elicitors on the accumulation of tanshinones in hairy root cultures of *Salvia miltiorrhiza. C. globosum* D38 mainly colonized in the intercellular gap of xylem parenchyma cells of *S. miltiorrhiza* hairy root, during long term co-existence without any toxicity against *S. miltiorrhiza* hairy root. Moreover, both of the live fungus and its mycelia extracts could induce the production of tanshinones, in special dihydrotanshinone I and cryptotanshinone. The effects of mycelia extracts were much stronger than that of the live fungus on tanshinones synthesis, which increased the transcriptional activity of genes with repect to tanshinone biosynthetic pathway obviously. Our results indicated that both of the live *C. globosum* D38 and its mycelia extracts could be utilized for tanshinones accumulation in *S. miltiorrhiza* hairy root. What’s more, D38 also could be made into biotic fertilizer applying into *S.miltiorrhiza* seddlings, which not only promoted host growth but the tanshinones and phenylpropionic acid accumulation. In the soil environment, D38 had formed bitrophic and mutual beneficial relationship with the host and enhanced the primary metabolism on the whole so as to have facilitative effects on phenylpropionic acid accumulation. To sum up, *Chaetomium globosum* D38 was a highly effective endophytic fungus for *S. miltiorrhiza*.

## Introduction

It was important to study host-microbe interactions, medicine or agriculture and especially for the influence of long-term microbial beneficially colonization on herbal medicine(Guo et al. 2015). Herbal medicines consumption has been increasing steadily and plant-derived secondary metabolites have become an important part for human health and nutrition. However, the yield of secondary metabolites in plants is rather little and depends greatly on physiological stages of the plants (Chandra and Chandra 2011). In recent years, cultured plant tissues have become a promising alternative source for bioactive secondary metabolites increasingly desired in medical and pharmaceutical field (Ahlawat et al. 2014). It is known that elicitation can enhance biosynthesis of secondary metabolites and biotic elicitors (Thwe et al. 2016) have particularly been used to induce the secondary metabolites accumulation (Bahabadi et al. 2014). It is one of the most effective means to induce secondary metabolites biosynthetic pathways in hairy roots or plant cells using pathogenic and nonpathogenic fungal as biotic elicitors (Huang et al. 2016). Generally, fungi can easily switch from an endophytic to necrotrophic lifestyle at the evolutionary and even ecological timescale. Different from necrotrophic fungal, the endophytes could infect host tissues with no concurrent symptoms of disease (Krings et al. 2007). In many cases, endophytes form mutualistic interactions and beneficial relationship with their host(Hyde et al. 2008, Rodriguez et al. 2009), which can not only stimulate plant growth, but also promote secondary metabolites accumulation in the plant (Murthy et al. 2014).

Plant tissue cultures, as the most convenient and useful experimental measures, are usually used to test various factors on desired products biosynthesis and explore effective measures to promote their production (Jian et al. 2005, Kai et al. 2011). However, not all the fungal endophytes have the ability to co-culture with their host plant tissue for a long time, which depends on the toxicity of the fungal isolate. In most cases, those fungal were fabricated into elicitors by the way of removing their toxicity, which can also enhance the secondary metabolites biosynthesis (Giauque and Hawkes 2013).

*Salvia miltiorrhiza* Bunge (Lamiaceae) is a famous and very important medicinal plant in China and the roots have been traditionally utilized to treat menstrual, cardiovascular and various inflammation-related diseases (Han et al. 2008, Wu and Yeung 2010). Tanshinones were a kind of bioactive compounds in *S. miltiorrhiza* roots, which demonstrates versatile pharmacological activities including antioxidant, cardiovascular protective, antibacterial, anti-inflammatory and antineoplastic activities (Tu et al. 2012, Koji et al. 2015). However, the low yield of tanshinones usually requests the use of large amount of plant material and thus appears as a major obstacle for *S. miltiorrhiza* exploit (Kai et al. 2012).

Treatment with elicitor is one of the most effective means for stimulating secondary metabolism in medicinal plants (Huang et al. 2016). It is reported that there have been few studies documented on the effects of elicitors, yeast extract (YE), salicylic acid (SA), and methyl jasmonate (MJ) on the tanshinones metabolism in *S. miltiorrhiza* hairy root (Hui et al. 2001, Ge and Wu 2005). Only our group has previously reported an elicitor from fungal endophyte *Trichoderma atroviride* D16, which could significantly promote the biosynthesis of tanshinone constituents (Ming et al. 2013). However, the hairy roots of *S. miltiorrhiza* can not be long-term co-cultured with *T. atroviride* D16. So we continued to search for other favorable endophytes, which could induce tanshinones accumulation without any toxicity against *S. miltiorrhiza* hairy root during long-term co-culture. This is therefore the first research on the effect of live endophytic fungus and its elicitor on the tanshinones production in *S. miltiorrhiza* hairy root. In this study, the tanshinones-promoting endophyte was identified as *Chaetomium globosum* D38. The effects of *C. globosum* D38 and its extract of mycelium (EM) on the tanshinones accumulation in *S. miltiorrhiza* hairy root cultures were further studied and the possible mechanism was also discussed for understanding the role of *C. globosum* D38 (Reissinger et al. 2003) in host survival. And the *Chaetomium globosum* D38, as a biotic fertilizer, had the same effects on *S. miltiorrhiza* seedlings, laying the foundation for practical applications.

## Materials and methods

### Isolation and identification of endophytic fungusn D38

First of all, in order to remove soil and dirt, the roots of *S. miltiorrhiza* were washed by running water, followed by deionized water. According to the literature, (Huang et al. 2009, Tan et al. 2012) the root was cut into 0.5 cm section and sterilized successively by 75% ethanol for 30 s, 1 min in 2.5% sodium hypochlorite, and 30 s in 75% ethanol. Then, they were flashed by sterile water for four times and desiccated by sterile filter paper. Finally, the tissues were placed and cultured on PDA medium at 28 °C,which contained 100 mg L^−1^ penicillin. The new hyphal was transfered separately on new medium and incubated subsequencely at 28 °C for 14 days till obtaining a pure strian.

The endophytic fungus was cultured on PDA medium at 28 °C for 7 days and was characterized by morphological features. The mycelium was scraped and its genomic DNA was extracted by using the CTAB method from the surface of the PDA medium. The ITS regions and 5.8 S gene were amplified by using the universal primers ITS5 and ITS4 (Min 2013) and compared by Blast search at the GenBank and aligned with CLUSTAL X software by using 1000 bootstrap replicates (Larkin et al. 2007). The phylogenetic tree was performed using the neighbor-joining method and identification at species taxonomic levels was based on ≥ 99% ITS similarity (Achille et al. 2006). The GenBank accession number of the nucleotide sequence was MF461354 and the endophytic fungus D38 has been kept in China General Microbiological Culture Collection Center in Beijing, China (collection number CGMCC 12379).

### Hairy root culture

*S. miltiorrhiza* hairy root was vaccinated to 250 ml conical flasks which contained 100 mL 1/2 B5 medium, which were then cultured at 25 °C, 135 rpm in darkness for 3 weeks. The medium was changed once every week and hairy roots were randomly divided into blank groups and experimental groups. The plaque was inoculated with a diameter of 0.5 mm inoculation loops to conical flask as experimental group and the same size of medium patch was inoculated into the blank groups. Endophytic fungi D38 was cultured with hairy root for 18 days and sampled on 0, 6, 12 and 18 day respectively. Each treatment has three repeats at each time for blank groups and experimental groups and nutrient solution was changed every 6 days.

### Extract of mycelium preparation and induction

Endophytic fungi D38 was inoculated to 250 mL conical flask by inoculating loop (5 mm in diameter). The conical flasks contained about 100 mL 1/2 B5 medium, and were cultured at 25 °C, 135 rpm for growing in the table concentrator. After 7 days, D38 was decompressed and filtrated, with D38 mycelium retained and washed 5 times by distilled water. After homogenated for 5 min, D38 mycelium solution was ultrasonically extracted for 60 min and then treated at 121 °C for 30 min in high pressure steam sterilization pot to remove free protein substances. Afterwards, the D38 mycelium solution was decompressed and filtrated as D38 elicitor.

### Immunocytochemical staining for the observation of D38 infection

Immunocytochemical staining procedure of root tissue was originally described by K. F. Störkuhl (Störkuhl et al. 1994). Fresh *Salvia miltiorrhiza* hairy root tissue was fixed in 4% paraformaldehyde for more than 24 h. The hairy root tissue was subsequently repaired level in the fume hood with a scalpel and then the tissue and the corresponding label were put in the dehydration box. Next, the dehydration box was put into the basket and dehydrated with gradient alcohol in turn. After that, waxed tissue was placed in the embedding machine and added with corresponding modification. Finally, waxed tissue was sliced in paraffin section by 4 μm. The slice was put into dimethylbenzene xylene I for 15 min, dimethylbenzene xylene II for 15 min, ethanol I for 5 min, anhydrous ethanol II for 5 min, 75% alcohol for 5 min, 85% alcohol for 5 min, and distilled water in turn. And then, the slice was put into the repair box which was filled with EDTA antigen repair buffer (pH 8.0) (Google biological technology co., Art.No.G1202). The antigen repair was performed in the microwave. After the medium was heated to boil for 5 min, cut off electricity for 5 min at interval. After natural cooling, the slice was placed into the PBS (pH 7.4) and washed on the decoloring table for three times, 5 min each time. After spin-dry, the slice was drew a circle with Pap Pen to prevent the antibody from running away. Dropping 3% BSA (solarbio, Art.No.A8020) in the circle, the tissue was covered uniformly and closed for 30 min at room temperature. The slice was placed flat on the wet box with some water and incubated for 24h at 4 °C after added with primary antibody (1:100 sword bean protein A with fluorescence). Then the slice was put into PBS (pH 7.4) for decoloration (washing three times, 5 min each time). After added with the corresponding second antibody in the circle, the slice was incubated for 50 min at room temperature in dark and then washed in the decoloration box for three times, 5 min each time. After spin-dry, the slice was added with DAPI (Google biological technology co., Art.No.G1012) and incubated for 10 min at room temperature in dark. After washed for three times and sealed with anti- fluorescence quenching tablets (Google biological technology co., Art.No.G1401), slice was inverted in the fluorescence microscope to collect images (Nasr et al. 2006).

### Manufacture of D38 fertilizer

There are 250 g wheat bran, 250 g cottonseed shell, 20 g glucose, 3 g KH_2_PO_4_, 1.5 g MgSO_4_ · 7 H_2_O dissolving in every 1 liter deionized water through sterilization for cultivating endophytic fungi D38. After fostering for 14 days, the number of spores was determined as 1~2 × 10^9^ by hemacytometry method. Subsequently, the 20g D38 fertilizers were applied into each *Salvia miltiorrhiza* aseptic seedling cultivated by sterile soil with sterile water whereas the blank was add by 20g blank wheat bran - cottonseed medium in July, 2016. And the cultured potted soil matrix is made of pearl cotton: humus: vermiculite with the ratio of 1:3: l. Each treatment has five repeats and was taken sampling every 2 months from September, 2016 to March, 2017 for the measurement of morphological indexes, biomass and bioactive substances content.

### HPLC analysis

*S. miltiorrhiza* hairy roots were desiccated at 55 °C in the oven and grinded into powder. After screened by a 40 mesh, hairy root powder was weighed 0.2 g accurately and put into 10 mL centrifuge tube. Accurately draw 4 mL of methanol using 5 mL pipette and add it into the centrifugal tube. After ultrasonically extracted for 60 min, the sample solution was filtered with 0.45 syringe filter, which was further used for HPLC analysis. The analysis was performed on an Agilent-1100 system using a ZOBAX-EXTEND-C18 chromatographic column (250 mm × 4.6 mm, 5 μm) at 30 °C with a H_2_O (+0.1% HCOOH) (A)/ acetonitrile (B) gradient (0-15 min: 80% B, 15-16 min: 80% B-62% B, 16-25 min: 62% B) as previously described(Ming et al. 2013). Rosmarinic acid, salvianolic acid B, dihydrotanshinone I, tanshinnone I, cryptotanshinone, and tanshinone IIA in the methanol extract of hairy roots were identified in comparison with the available standards. The reference standards were aquired from Chengdu Mansite Pharmacetical Co. Ltd. [Chengdu (Sichuan Province), PR China].

### RNA isolation and real-time quantitative PCR analysis

Hairy roots were divided randomly into blank group and D38 groups which were cultured for 18 days and nutrient solution was changed every 6 days. Hairy roots of two groups were sampled at 0, 6, 12, and 18 day and preserved in –80 °C refrigerator. The total RNA of samples were extracted with mir Vana™ miRNA isolation Kit (TaKaRa). After detecting the good integrity of total RNA and obtaining cDNA by reversed transcription, the reaction was started on fluorescence quantitative PCR instrument (LightCycler II 480). The list of primers used in the real-time PCR reaction is signed as table1 for detecting the gene expressions of *smHMGR, smDXR, smGGPPS, smCPS*, and *smKSL*, respectively (Ma et al. 2012). The 18S gene was used as a reference gene to normalize cDNA. The process of RT-PCR reaction is set as follows: initial degeneration at 95 °C for 15 min, subsequently 40 cycles of 95 °C degeneration for 10 s, and finally 60 °C annealing extending for 30 s. The specificity of product was detected by using the melting curve, in which temperature was elevated slowly from 60 °C to 97 °C, gathering five fluorescent images every °C. The relative gene expression was quantified using the comparative CT method.

**Table1.**
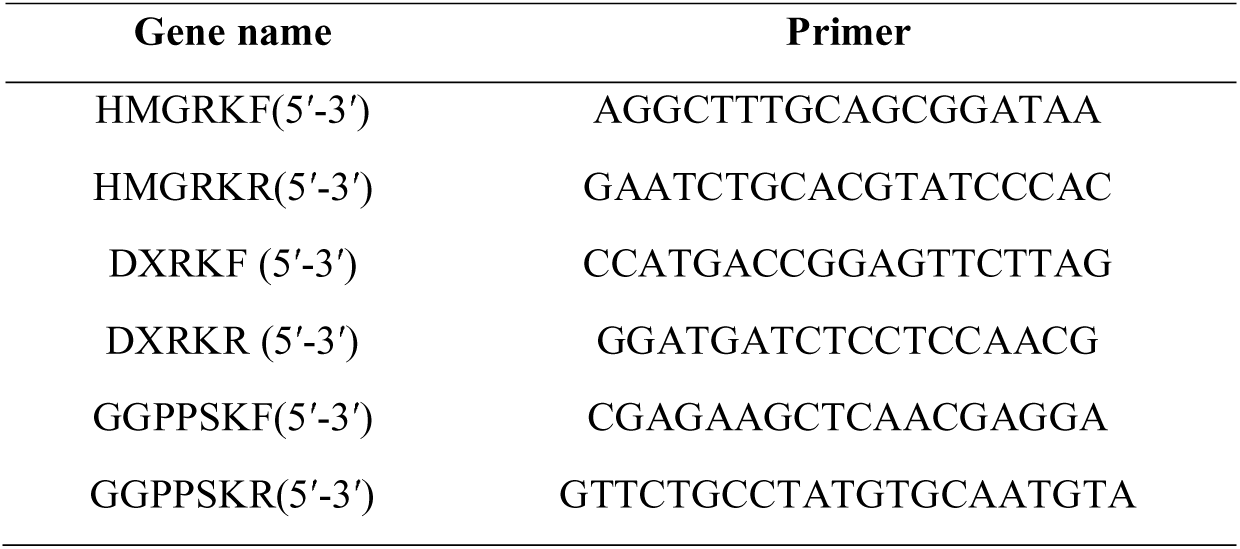
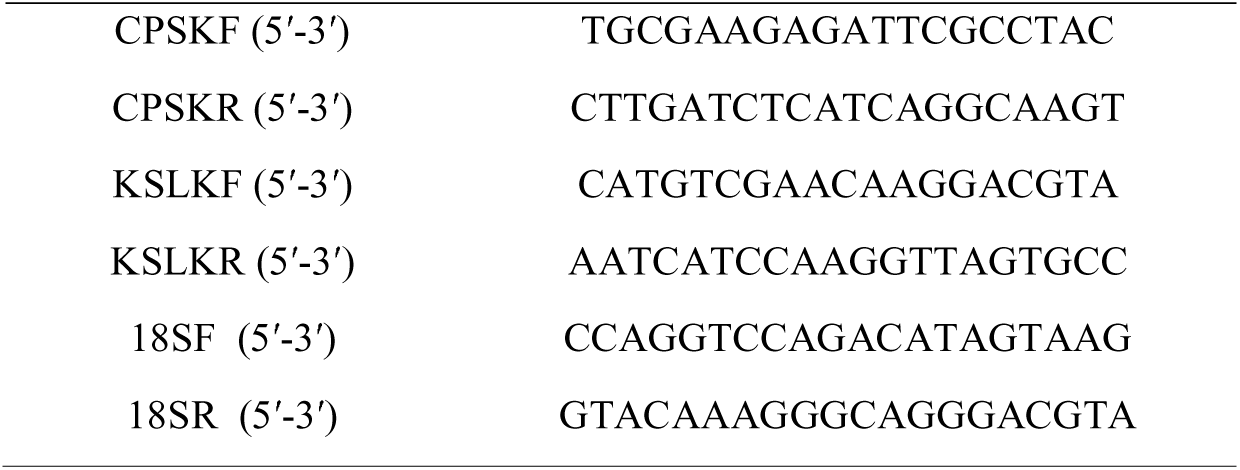
The real time-PCR primer of gene.

### Data analysis

The data of HPLC analysis and semi-quantitative real-time PCR of hairy root cultures were anlyzed by one-way analysis of variance (ANOVA) through SPASS software which contained both control and different treatments in triplicate. The results are presented as mean ± SD (standard deviation) and the error bars represent the standard deviation of biological triplicates in the figures. Otherwise, the term significant has been used to denote the differences for which *p* is <0.05 and the statistical significance of differences in gene transcripts was analyzed by one-sample *t*-test.

## Results

### Isolation and identification of endophytic fungus D38

The D38 on solid culture is characterized by surface fleece with similar yellowish color on the back, and microstructure of D38 is long chain without branches (Figure 1). In this study, the internal transcribed spacer region (ITS) of D38 was selected and sequenced with nuclear ribosomal DNA as a template. Results of the BLAST searches for D38 strain showed that the ITS rDNA sequences of the isolate shared high homology with *Chaetomium globosum*. The sequence identity of the ITS and 5.8S was 99% (Figure 2). According to the molecular data, the strain was identified to the genus *Chaetomium globosum*.

**Figure 1.**
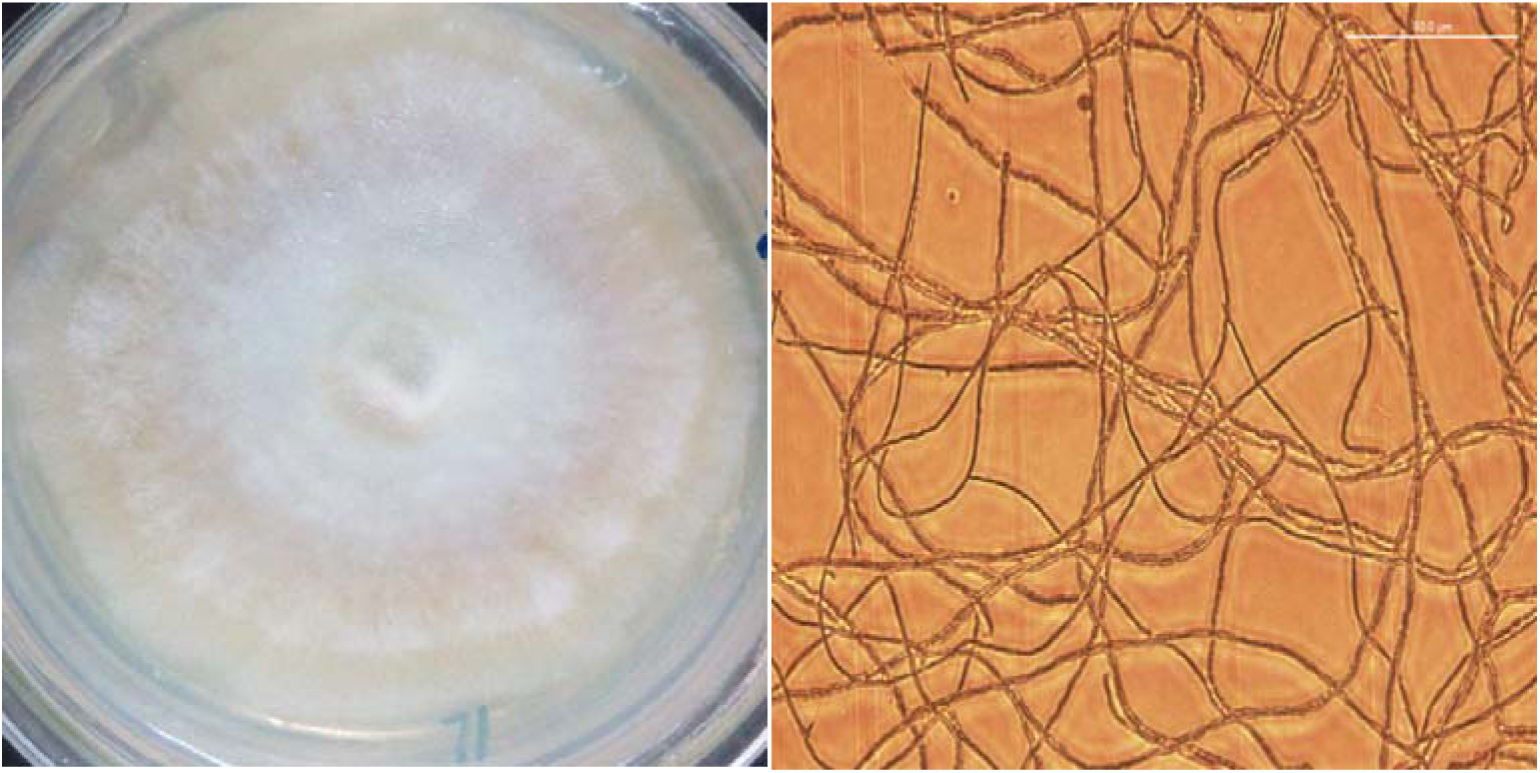
The morphological characteristics and microstructure of D38 (A) Colony positive photo; (B) Micrograph of 3R-2.

**Figure 2.**
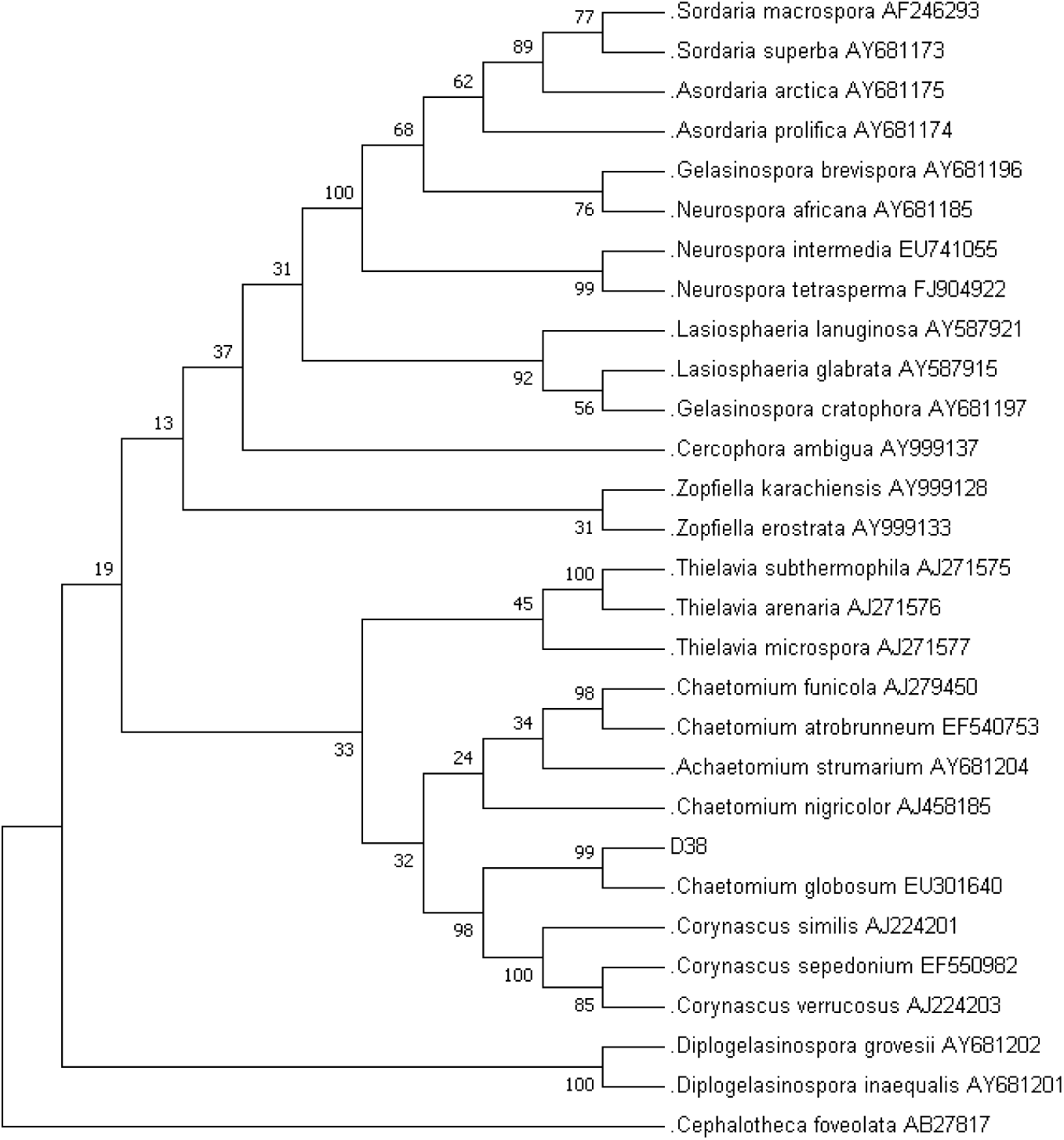
The phylogenetic tree of the endophytic fungus D38, Cephalotheca foveolata (AB27817) is used as outgrout.

### Effects of D38 on the contents of tanshinones in S. miltiorrhiza hairy roots

During our preliminary study, endophytic isolate D38 can be long-term co-cultured with *S. miltiorrhiza* hairy roots. Further we investigated the effects of the live fungus D38 on tanshinones biosynthesis in *S. miltiorrhiza* hairy roots. As shown in Figure 1, administration of D38 significantly enhanced the contents of dihydrotanshinone I (DT-I) and cryptotanshinone (CT) on day 6, 12 and 18, compared with that of blank group. The content of dihydrotanshinone I and cryptotanshinone were increased by 9 times and 13.2 times, respectively, on day 18 (Figure 3a and 3b). However, no such notable activity were observed for D38 on the contents of tanshinone I (T-I) and tanshinone IIA (T-IIA). D38 only increased the content of tanshinone I on day 6 (1.8 time) (Figure 3c) and elevated the content of tanshinone II A on day 6 (1.7 time) and 18 (1.9 time) (Figure 3d).

**Figure 3.**
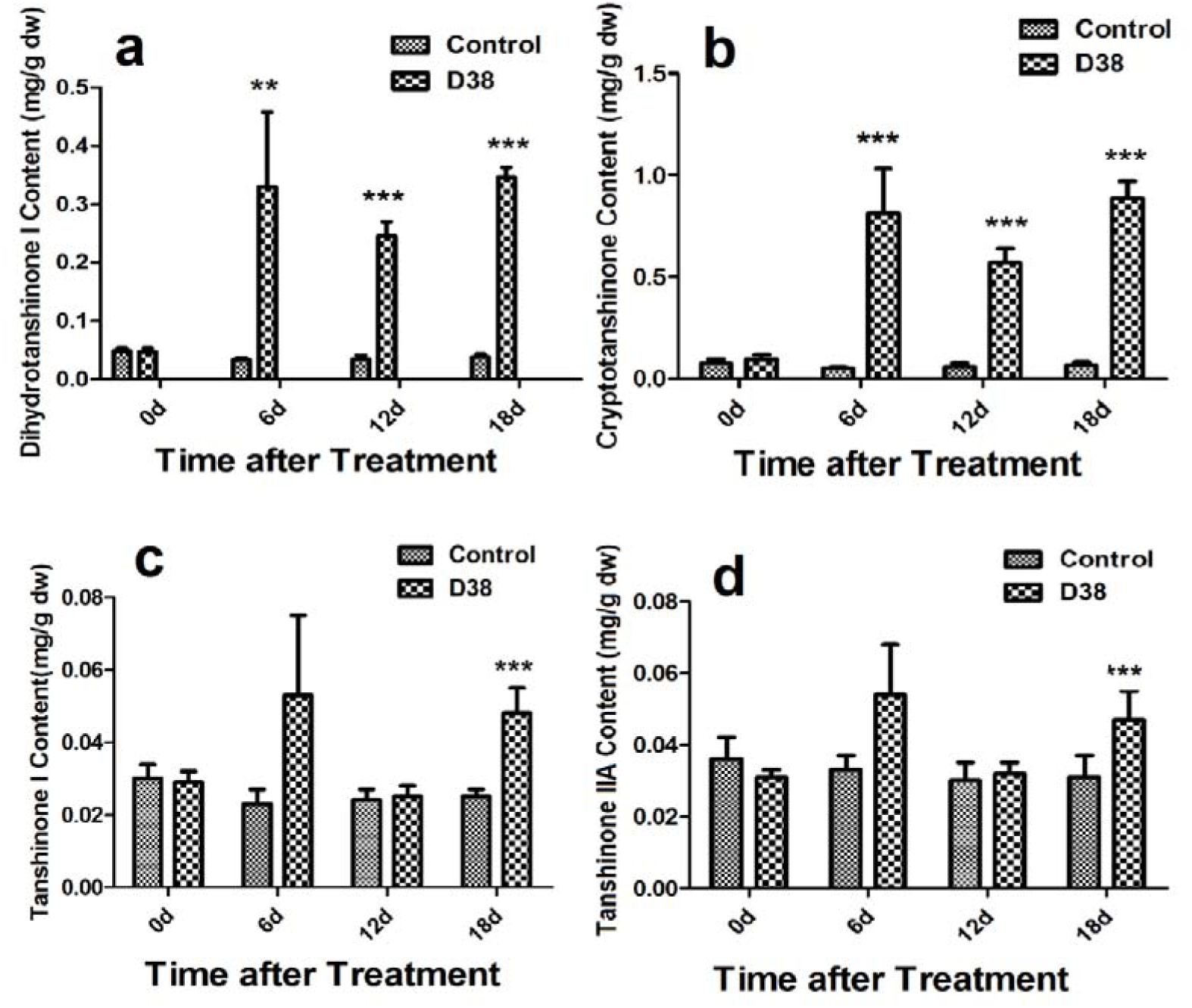
The effect of D38 hyphae on the contents of tanshinones of *S. miltiorrhiza* hairy roots.Values are presented as means ±SD, n=3. *P < 0.05; **P < 0.01; ***P < 0.001 versus the control culture.

### Immunocytochemical staining of D38 in S. miltiorrhiza hairy roots

After the co-cultivation of *S. miltiorrhiza* hairy root with endophytic fungi D38 after 18 days, the immunofluorescence staining experiment was performed to display the infection location of D38 (Figure 4). Endophytic fungi D38 infected the cell gap of S. *miltiorrhiza* hairy root tissue, and minority of D38 hypha existed in the tissue cells. Thus we speculated that mycelium of endophytic fungi D38 started invading into the internal cells and form the enveloped between intercellular space, which manisfied that D38 could interact with host stably. In conclusion, we found that endophytic fungi D38 mainly colonized in the intercellular gap of xylem parenchyma cells of *S. miltiorrhiza* hairy root as a beneficial endophyte of *S.miltiorrhiza*.

**Figure 4.**
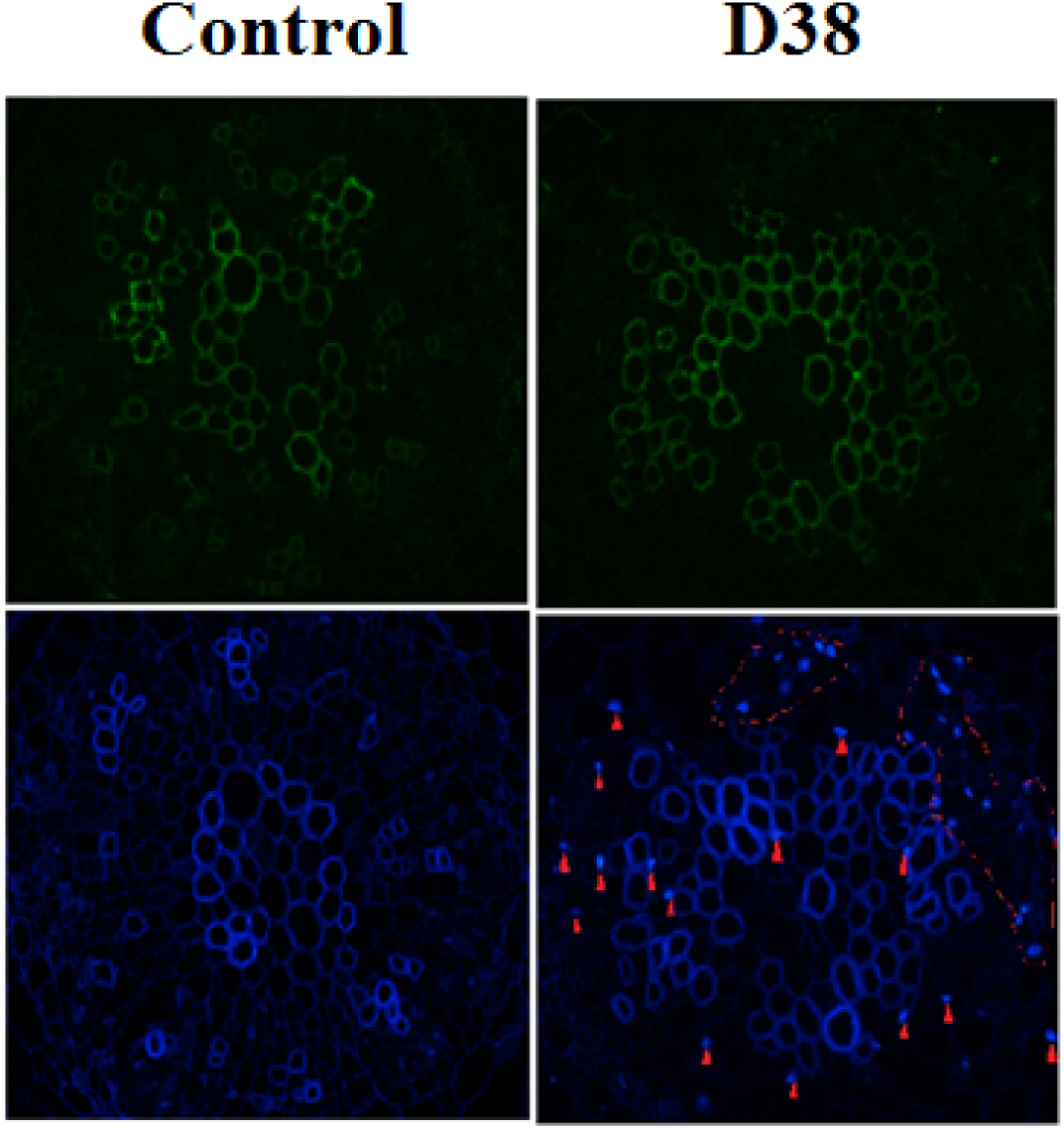
The endophytic fungus D38 in *S. miltiorrhiza* hairy roots after immunofluorescence staining (the red arrows and the circles indicates the mycelia, amplification factor : 400×)

### Effects of extract of D38 mycelium (EM) on the tanshinones accumulation in S. miltiorrhiza hairy roots

After the treatment of co-culture between hairy root and different concentration of D38 EM, *S. miltiorrhiza* hairy roots were then extracted ultrasonically with methanol for HPLC analysis. The data revealed content of tanshinones increased significantly under the co-culture with D38 inducer solution (Figure 5). And the content of tanshinone I and cryptotanshinone increased most significantly. On the 18^th^ day, the content of dihydrotanshinone I and cryptotanshinone increased most significantly. The content of dihydrotanshinone I reached the highest under the action of the 60 mg/L D38 EM and improved by 22 times compared with the blank group. However, the content of cryptotanshinone reached the highest under the action of 90 mg/L D38 EM and increased by 20.3 times compared with blank group. At the same time, tanshinone I and tanshinone IIA had obvious enhancement under the action of different concentration of D38 EM. The content of tanshinone I reached the highest by 2.2 times higher than that of blank group under 90 mg/L D38 EM and the content of tanshinone IIA reached the highest by 2 times compared with the blank group under the action of 90 mg/L D38 EM.

**Figure 5.**
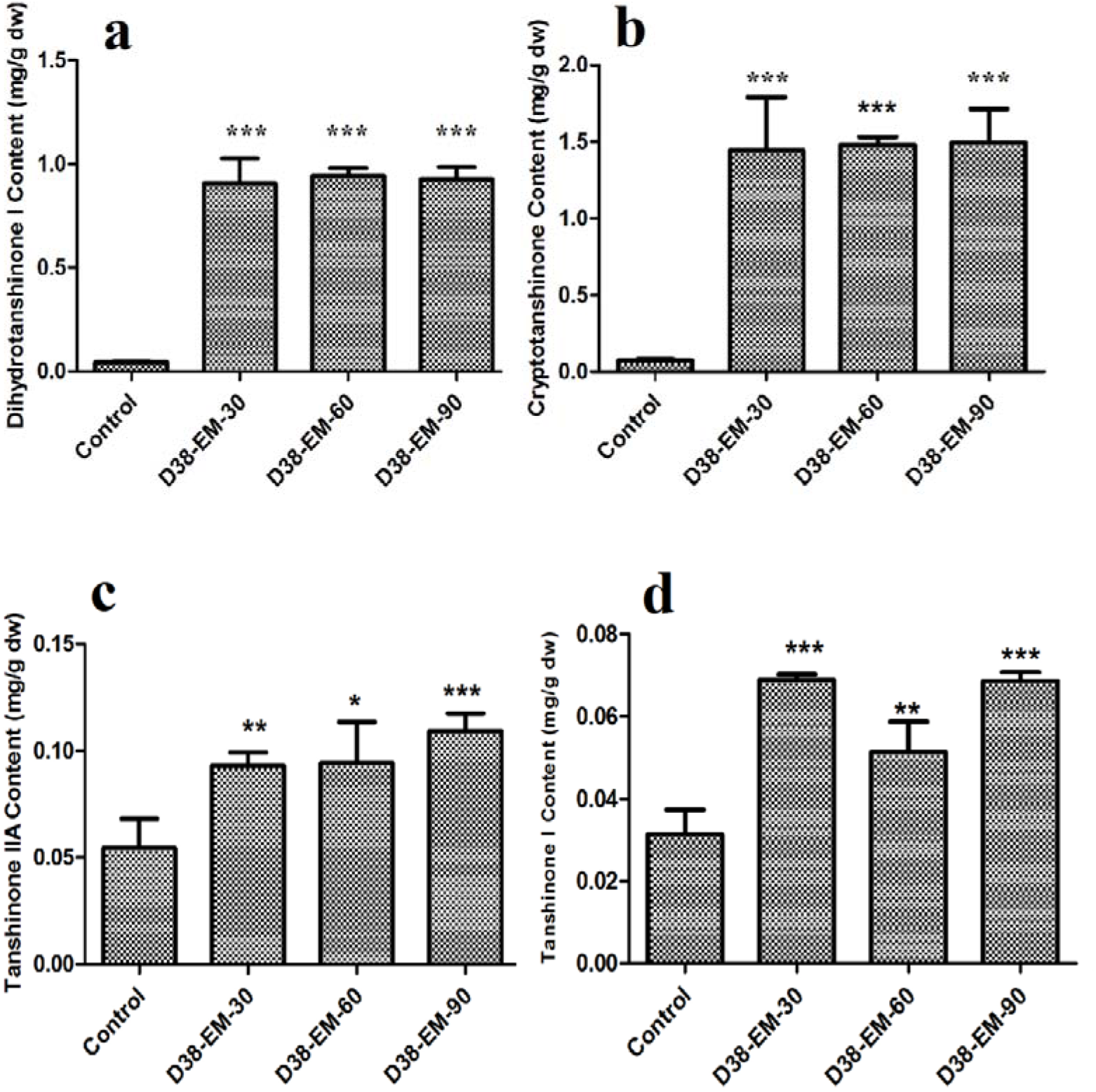
Effects of the extract from D38 mycelium (EM) on the contents of tanshinones in *S. miltiorrhiza* hairy roots.Values are presented as means ±SD, n=3. *P < 0.05; **P < 0.01; ***P < 0.001 versus the control culture.

### Transcriptional response of the tanshinone biosynthetic pathway to the extract of D38 mycelium in S. miltiorrhiza hairy roots

After extracting RNA and get cDNA through the reverse transcription reaction, the transcription level of five key enzyme genes were determined by real-time quantitative PCR in the tanshinone biosynthesis pathway (Figure 6).

**Figure 6.**
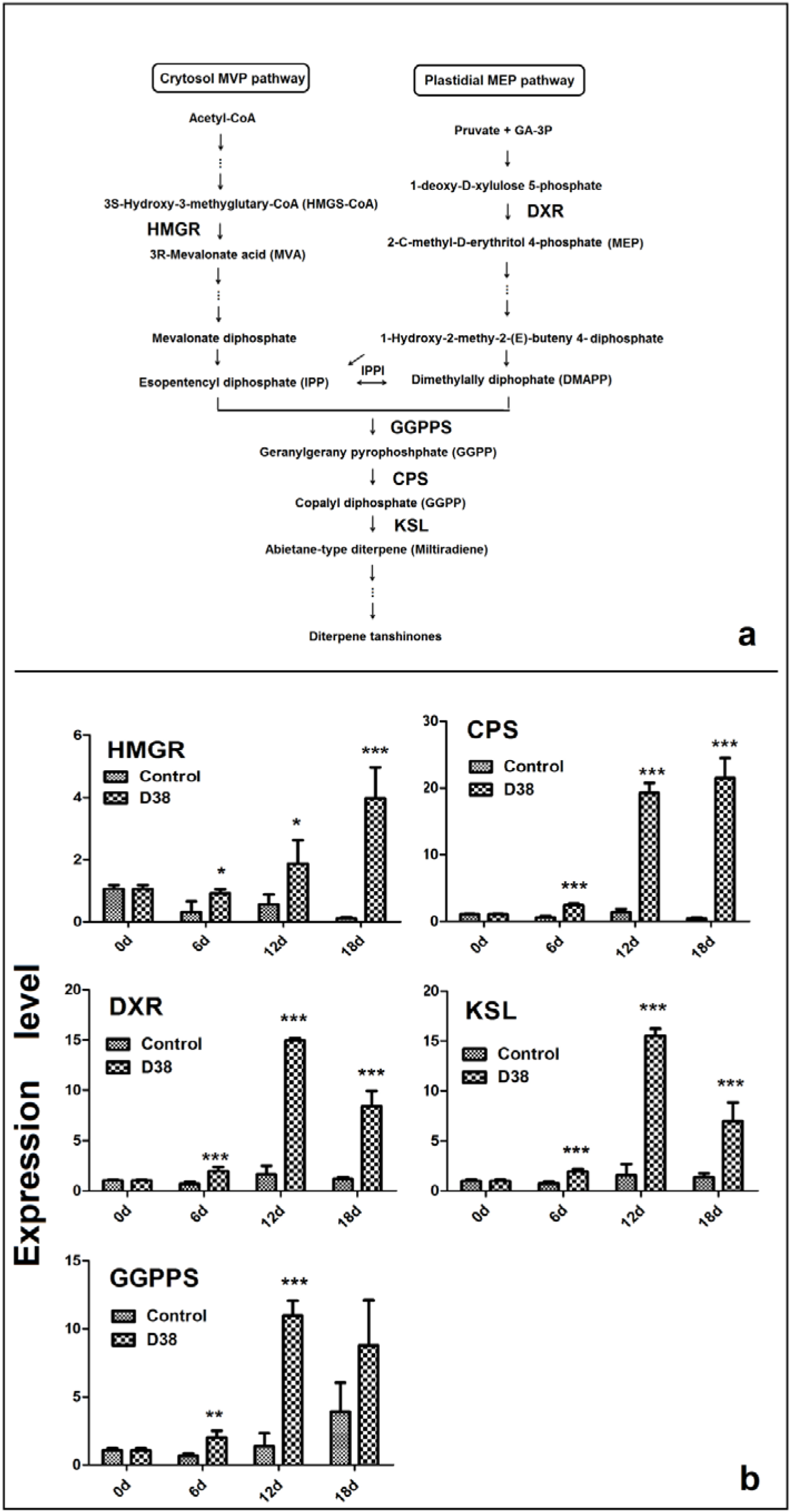
Proposed pathways of tanshinone biosynthesis in *S. miltiorrhiza* (A) and effects of the D38 EM (100 mg·l^−1^) on the expression of genes in the tanshinone biosynthetic pathway in *S. miltiorrhiza* hairy roots in 18^th^ day (B). HMGR, hydroxymethylglutaryl-CoA reductase; DXR, 1-deoxy-d-xylulose 5-phosphate reductoisomerase; GGPPS, geranylgeranyl diphosphate synthase; CPS, copalyl diphosphate synthase; KSL, kaurene synthase-like. Values are presented as means ±SD, n=3. *P < 0.05; **P < 0.01; ***P < 0.001 versus the control culture.

D38 EM activated the five key enzymes in the synthesis pathway and the levels of key enzyme genes expression have significantly improved. The expression level of 3-hydroxy-3-methylglutaryl-CoA reductase (HMGR) reached highest by 36 times in 18^th^ day compared with the blank group invovled in the mevalonate (MVA) pathway.In the 2-C-methyl-D-erythritol-4-phosphate (MEP) pathway, the expression level of 1-deoxyd-xylulose 5-phosphate reductoisomerase (DXR) reached the highest by 9.35 times compared with the blank group in 12^th^ day. In the downstream pathway, the expression level of geranylgeranyl diphosphate synthase (GGPPS), copalyl diphosphate synthase (CPS), kaurene synthase-like (KSL) was enhanced significantly, especially the 12^th^ and 18^th^ day. The expression level of GGPPS and KSL reached highest by 8 times and 10 times respectively in 12^th^ day. The expression level of CPS reached the highest expression level in 18^th^ day by 55.1 times compared with the blank group.

### Effects of D38 fertilizer on the tanshinones accumulation in S. miltiorrhiza roots

After the treatment of co-culture between each *S.miltiorrhiza* seedling and 20g D38 fertilizer for half one year from July, 2016 to December, 2016, the morphology indexes of *S. miltiorrhiza* were recorded including the numbers of leaves, plant height, wet weight and dry weight. At the same time, *S.miltiorrhiza* roots were then extracted ultrasonically with methanol for HPLC analysis every two months. The data revealed D38 fertilizer can promote S.miltoirrhiza growth greatly (Supplemental Fig.1) and the content of salvianolic acid B and tanshinones increased significantly under the co-culture with D38 (Figure 7,8). And the number of leaves, plant height, wet weight and dry weight is keeping increased. The number of leaves was increased under the treatment of D38 fertilizer in Sept. 2016, however, the leaves were growing but some of leaves were fallen. Therefore, the number of leaves was decreased in Nov. 2016 compared with Sept. 2016. And the plant height of D38 fertilizer was larger than blank group. At the same time, the wet weight and dry weight was 1.95 fold and 5.20 fold than the control in Sept. 2016, and kept the rise rate in the following two months. That was to say D38 was a growth-promoting endophyte, which can facilitate the growth of *S.miltoirrhiza* hairy root.

**Figure 7.**
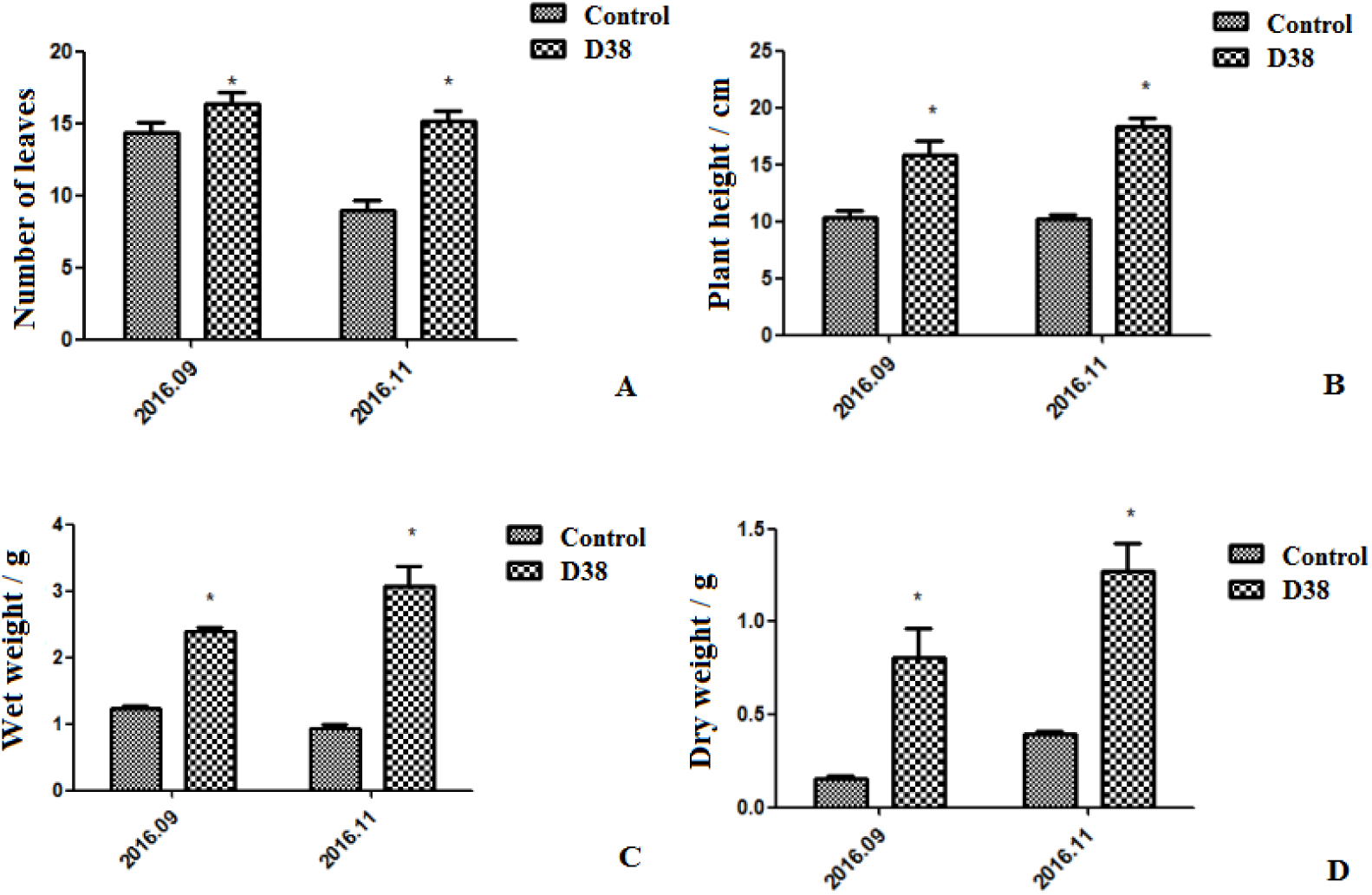
The measurement of morphological indexes under the treatment of 20g D38 fertilizer compared with blank group from September, 2016 to December, 2016. (A) Number of leaves; (B) Plant height; (C)wet weight; (D) Dry weight.

**Figure 8.**
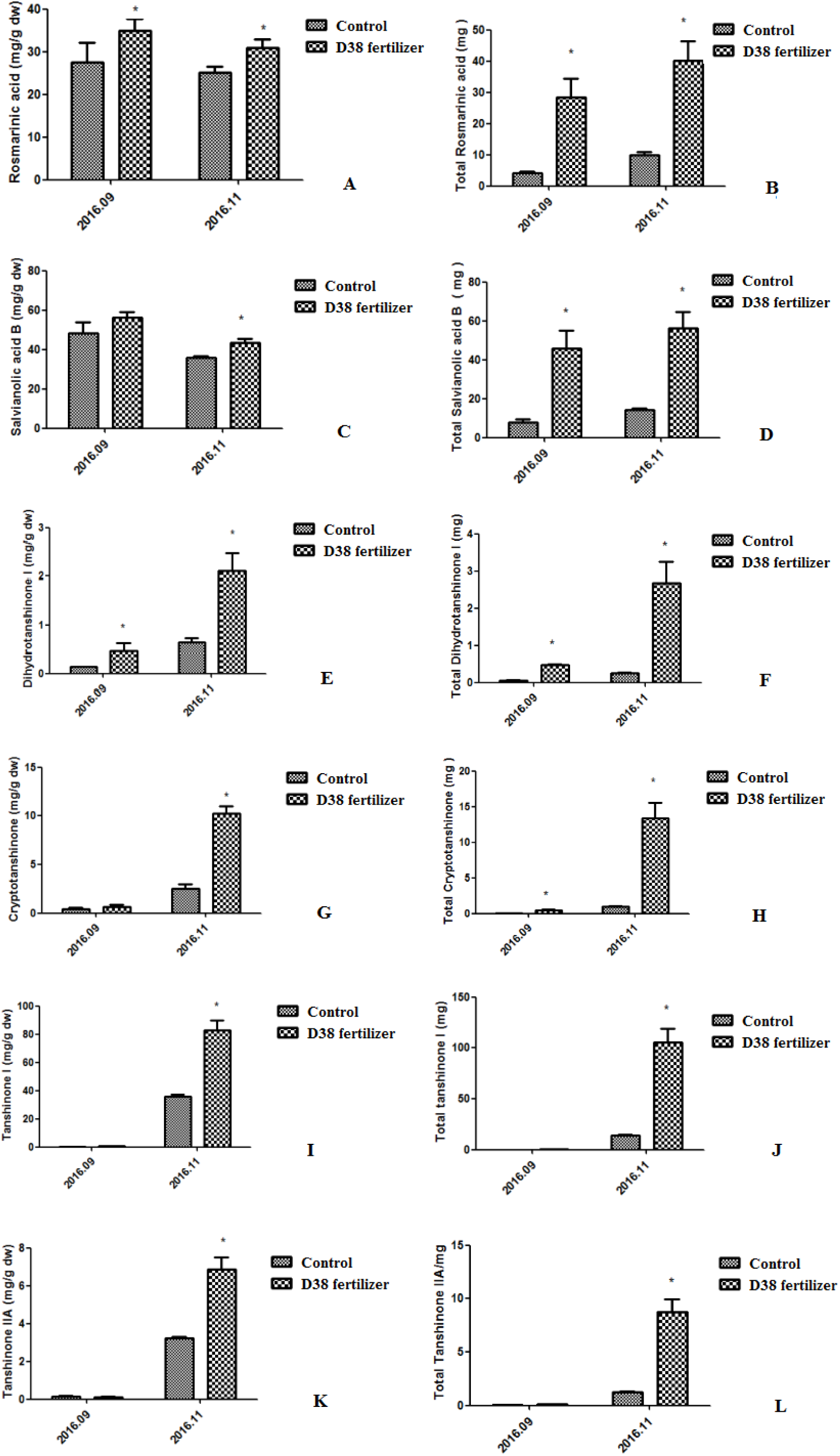
The measurement of bioactive substances of *S.miltiorrhiza* root under the treatment of 20g D38 fertilizer compared with blank group from September, 2016 to December, 2016. (A) Rosmarinic content per unit of root mass; (B) Total Rosmarinic content; (C) Salvianolic acid B content per unit of root mass; (D) Total Salvianolic acid B content; (E) Dihydrotanshinone I content per unit of root mass; (F) Total Dihydrotanshinone I content; (G) Cryptotanshinone per unit of root mass; (H) Total Cryptotanshinone content; (I) Tanshinone I content per unit of root mass; (J) Total Tanshinone I content; (K) Tanshinone IIA content per unit of root mass; (L) Total Tanshinone IIA content.

In addition, phenolic acid and tanshinones content per unit of root mass was enhanced drastically under the treatment of D38 fertilizer in November, 2016 and the total content of bioactive substances is increasing continually during the half year. The rosmarinic acid content per unit of D38 fertilizer treatment was higher than the control by 27% and 24% in Sept. and Nov. 2016 separately. The salvianolic acid B content per unit of D38 fertilizer treatment was higher than the control by 21% in Nov. 2016. And a significant 22% boost of salvianolic acid came up in Nov. 2016. And the contents of dihydrotanshinone I, cryptotanshinone, tanshinone I and tanshinone IIA per unit were dramatically enhanced by 2.29 fold, 3.07 fold, 1.30 fold and 1.14fold in Nov. 2016. After 5 months growth, the root of *S.miltoirrhiza* has entered therapid growth period and the effects of D38 fertilizer was becoming more significant, which indicated D38 fertilizer could be developed into new biotic fertilizer.

## Discussion

Plants were associated with various of microbes, containing pathogens, mycorrhizal fungi, rhizosphere bacteria, and endophytes (Crawford et al. 2010). Most plants infested by fungi caused no disease symptoms in natural ecosystems. Due the bioactive compounds belonging to secondary metabolites are usually shorter than primary metabolites in medicinal plants, a complex network of reactions could be triggered by all kinds of fungal, which ultimately cuase the synthesis and accumulation of secondary metabolites(Zhao et al. 2010). For example, an endophytic fungus was found in the bark of the *Taxus chinensis* tree and could produced three times as much taxol as non elicited cells (Wang et al. 2001). There are few endophytes isolated from the host plant which can co-culture with the host plant tissue for a long time, however, fungal endophytes act as a zoetic elicitor that can multiply and persistently stimulate the plant tissue(Saikkonen et al. 2004, Rusch 2016). The effects of plant-microbial interactions depend on the biological and non-biological factors and the genotypes and interactions of the host(Brader G 2017). It was reported that successfully infected endophytes may overcome the barrier of periderm of root and enter the above-ground of plant via central cylinder(Sesma and Osbourn 2004, Sukno et al. 2008). Like some endophytes, D38 was also separated from the leaves of *S.miltiorrhiza,* which indicated D38 had form bitrophic interaction between D38 and host not only in root but leaves. Due to the process of endophyte infection, the hyphae may invade into epidemic cell and then develop the junction to form intercellular hyphae. After transient interaction, the intact host plasma membrane beseted intracellular hyphae just for the sake of longer interaction in cortical cells (Kei et al. 2016). Endophyte have a symbiotic relationship with plants, which may experience a long evolutionary and interacting communication (Wawra et al. 2016). The mycelium of endophytic fungi developed the best circle by colonizing in intercellular space, verifying a stable relationship between endophyte and host. In conclusion, endophytic fungi D38 finally reside in the intercellular gap of xylem parenchyma cells of *S. miltiorrhiza* hairy root as a beneficial endophyte of *S.miltiorrhiza* after a series of communication.

In this study, the accumulation of tanshinones in *S. miltiorrhiza* hairy roots was enhanced by both endophytic *Chaetomium globosum* D38 and its EM, which indicated D38 exhibited obvious effects on secondary metabolism of *S.miltiorrhiza* (Fig. 3, Fig. 5). EM is predicted to be one of the main active constituents responsible for promoting biosynthesis of tanshinones in *S. miltiorrhiza*hairy root cultures by comparing the effects of endophyte *Chaetomiumglobosum* with EM. In addition, the transcriptional levels of EM genes involved in the tanshinone biosynthesis pathway is increased obviously (Fig. 6). However, our present work is the first test to include *Salvia miltiorrhiza*hairy root co-cultured with endophytic fungal as an effective strategy for improving tanshinones production in *S Miltiorrhiza* hairy root. As far as we known, the effects of endophytic fungi and its elicitors on the secondary metabolism of their host plants was rarely reported., Fungal endophyte infection can change the metabolic profiles of *Lolium perenne*by carbon/nitrogen exchange (Rasmussen et al. 2008) and the active substances content of *S.miltiorrhiza,* was enhanced by beneficial endopyhtes D38, which predicted D38 and its host may has some form of substance communication.

One report indicated that tanshinones had much stronger antimicrobial activity than phenolic acids among which DT-I and CT exhibited the stronger antimicrobial effect. This may be as a potiential reason for EM, isolated from *Chaetomiumglobosum* D38 dramatically stimulating the biosynthesis of DT-I and CT (Fig. 5). The phenomenon interpreted into *S.miltiorrhiza* defending itself and responsed to the invading of *Chaetomiumglobosum* D38 mycelium through secreting more DT-I and CT when *Chaetomium globosum* D38 entered the *S. miltiorrhiza* hairy root.

Tanshinones were one kind of bioactive compounds as abietanoid diterpenes in *S. miltiorrhiza* roots, owning mighty and wide therapeutic effects. Nevertheless, the components account for a low proportion in *S.miltiorrhiza* and regulated by all kinds of rate-limiting enzymes (Fig. 6). Isopentenyl diphosphate (IPP) and dimethylallyl diphosphate (DMAPP) were general precursors of terpenoids in plants and synthesized via mevalonate (MVA) and 2-C-methyl-d-erythritol phosphate (MEP) pathway. MVA pathway proceeded in the cytoplasm while the other MEP pathway carried out in the plastids. HMGR is one of significant enzymes in the MVA pathway, while DXR in the MEP pathway. GGPPS catalyse was the junction enzyme between MVA and MEP pathway, indicating the significance of GGPPS. As to the downstream of tanshinone biosynthetic pathway, CPS and KSL are the key point as we know. For testing the effects of EM of D38, the transcript levels of HMGR, DXR, GGPPS, CPS, and KSL were investigated by real-time quantitative PCR (Fig. 4). From 6^th^ day to 12^th^ day, the transcriptional levels of the five genes were elevated significantly. The biosynthesis of tanshinones was evidenced mainly occuring via the MEP pathway, with dependence on the crosstalk between the MEP and the MVA pathways (Ming et al. 2013). Gene expression levels of GGPPS, CPS, and KSL were also observed to increase gradually with EM treatment while a little decline occurred in 18^th^ day. These results showed that EM stimulates many of the genes in the biosynthesis of tanshinones and then promotes the accumulation of tanshinones in S. miltiorrhiza hairy roots. However, it kept doubt how EM stimulated these genes through signal transduction pathway during the process, which need further study

In the present study, the results suggested that *Chaetomium globosum*D38 and EM from *Chaetomium globosum*D38 could elicit defence responses in the host plant as confronting pathogens. EM increased the secondary metabiolites of the host by enhancing the expression of related genes involved in the biosynthesis pathway. Therefore, EM can be used as a potent elicitor for stimulating tanshinone production in *S. miltiorrhiza* hairy root cultures. As a result, EM could act as a convenient strate for the extensive production of tanshinones in *S. miltiorrhi*za hairy root culture systems. The effects of EM are higher than the live endophytic fungal on the tanshinones synthesis, however, the treatment of live *Chaetomiumglobosum*D38 is successional and long-time means of stimulating the accumulation of plant tanshinones.Live *Chaetomium globosum* D38 is responsible for stimulating the biosynthesis of tanshinones in the hairy root culture and it is possible that live endophytic fungal can be effectively utilized for large-scale production of tanshinonesin the *S. miltiorrhiza* hairy root culture system in future studies, which may be greater and easier than fungal elicitor. At the same time, *Chaetomium globosum* D38 can promote the *S. miltiorrhiza* seedlings growth and the content of salvianolic acid B and tanshinones in *S. miltiorrhiza* root, which is a great breakthrough in practical *S. miltiorrhiza* cultivation. This could be interpreted that D38 can enhance the primary metabolism fluxes causing the increase of phenylpropionic acid content in soil environment. The number of leaves, plant height, wet weight and dry weight is keeping increased under the treatment of D38 fertilizer. As we all know, infection by the fungal endophyte may affect the accumulation of nutrition, such as inorganic and organic N (Bethlenfalvay et al. 1982, Lyons et al. 1990). In additon, D38 was also separated from the stem of *S.miltiorrhiza* and it may have transfered into the above-ground of plant by central cylinder. It signified that D38 has formed bitrophic interaction and reciprocity relationship in nutrients with *S.miltiorrhiza (Hacquard et al. 2016)*. Clonization of D38 in the host was a long selected and interacting process (Wawra et al. 2016), indicating D38 is a benefical endophyte for *S.miltiorrhiza*.

## Acknowledge

The authors declare that they have no conflict of interests. The work was supported by supported by the National High Technology Research and Development Program of China (No. 2014AA 020508) and the “Chen Guang” project supported by Shanghai Municipal Education Commission, Shanghai Education Development Foundation (13CG40) and National Nature Science Foundation of China (No. 81473301).

---

**Supplemental Figure 1.** The figure of under the treatment of 20g D38 fertilizer compared with blank group from September, 2016 to December, 2016. (A) The *S. miltiorrhiza* seedlings of blank group in September, 2016; (B) The *S. miltiorrhiza* seedlings of 20g D38 fertilizer in September, 2016; (C) The *S. miltiorrhiza* seedlings of blank group in December, 2016; (D) The *S. miltiorrhiza* seedlings of 20g D38 fertilizer in December, 2016.

